# A Syngeneic Immunocompetent Mouse Model of Gallbladder Cancer Reveals Tumorigenesis and Therapeutic Response

**DOI:** 10.1101/2025.11.17.687512

**Authors:** Wenqing Qiu, Xiaojian Ni, Fuqing Xiang, Weida Meng, Rongyi Qin, Jiakang Ma, Dandan Zhang, Bosen Li, Shijie Xu, Changcheng Wang, Lingxi Nan, Xiaobo Bo, Yun Liu, Li Wang, Yueqi Wang, Houbao Liu

**Affiliations:** Department of General Surgery, Central Laboratory, Shanghai Xuhui Central Hospital, Fudan University, Shanghai, China; Department of Biliary Surgery, Zhongshan Hospital, Fudan University, Shanghai, China; Biliary Tract Disease Institute, Fudan University, Shanghai, China; MOE Medical Basic Research Innovation Center for Gut Microbiota and Chronic Diseases, Wuxi School of Medicine, Jiangnan University, Wuxi, China; MOE Key Laboratory of Metabolism and Molecular Medicine, Department of Biochemistry and Molecular Biology, School of Basic Medical Sciences, Fudan University, Shanghai, China; Department of Gastrointestinal Surgery, Zhongshan Hospital, Fudan University, Shanghai, China; R&D Department, Suzhou Jiyan Biopharmaceutical Technology, TaicangBiomedical Industrial Park, Suzhou, China

## Abstract

**Purpose:** Gallbladder cancer (GBC) is a rare but highly lethal malignancy with one of the poorest prognoses among cancers of the digestive system cancers. However, current experimental models of GBC face critical limitations, particularly their dependence on immunodeficient hosts, which precludes the investigation of the tumor microenvironment. We aimed to establish a novel syngeneic immunocompetent mouse model that recapitulates human GBC and enables the investigation of tumor-immune interactions.

**Experimental Design:** We engineered a murine cell line (mGBC1-ZH) from normal mouse gallbladder organoids expressing *Kras* and *Trp53* (encoding mouse *p53*) mutations. Tumorigenic potential of mGBC1-ZH was evaluated by subcutaneous and orthotopic implantation. RNA-Seq and WES was used to demonstrate its characterization and similarity with human GBC. Immunohistochemistry, CCK8, and transwell assays wereused to investigate the role of CXCL5 in GBC. Therapeutic responses to standard first-line chemotherapeutic agents was evaluated in the syngeneic GBC model.

**Results:** This model supports both subcutaneous and orthotopic tumor growth in immunocompetent C57BL/6J hosts while preserving hallmark features of human GBC, including biliary epithelial differentiation (CK7+/CK19+), aggressive histopathology, and an immunosuppressive “cold” tumor microenvironment. Genomic characterization revealed recurrent chromosomal instability and copy number alterations mirroring human GBC. Comprehensive transcriptomic profiling revealed profound gene expression changes during the model development, and resembled transcriptional features between mGBC1-ZH and human GBC cell lines and samples. CXCL5 was found to be upregulated in human GBC, and promote proliferation, metastasis, and neutrophils infiltration of GBC in the syngeneic mouse model. Functional validation of the syngeneic GBC model demonstrated therapeutic responsiveness to frontline chemotherapeutics gemcitabine and cisplatin with significant in vivo tumor regression.

**Conclusions:** In summary, we establised a novel syngeneic GBC mouse model, overcoming the limitations of traditional models by enabling studies in immunocompetent hosts. This model provides valuable insights into the molecular evolution from normal gallbladder cells to transformed cancer cells and establishes a robust platform for both mechanistic studies and therapeutic development.

## Introduction

Gallbladder cancer (GBC) is a rare but highly lethal cancer originating from the mucosal layer of the gallbladder. Despite its rarity, GBC accounts for 80-95% of all biliary tract cancer and ranks sixth among gastrointestinal malignancies worldwide (1), carrying one of the poorest prognoses among digestive system cancers (2). The stark reality of GBC treatment is particularly concerning-while surgical resection offers the only path to long-term survival, fewer than 20% of patients qualify for this intervention (1). Even with recent advances in the combination of chemotherapy and immunotherapy, median survival remains low at 12.8 months (3), underscoring an urgent need for more effective therapeutic strategies.

The limited progress in GBC treatment stems largely from our incomplete understanding of its biological complexity. This knowledge gap persists despite significant advances in cancer biology, primarily owing to the lack of appropriate experimental models. Current approaches using established human GBC cell lines (4), patient-derived xenografts (PDXs) (5), and tumor organoids (6,7), although valuable, face critical limitations. Their dependence on immunodeficient hosts precludes the investigation of tumor-immune interactions - a crucial consideration given the emerging importance of immunotherapy. Moreover, these models, derived from end-stage malignancies, offer limit insight into the sequential events driving tumor development and progression.

Recent genomic analyses have revealed a complex molecular landscape in GBC, characterized by recurrent alterations in key tumor suppressors (*TP53*, *SMAD4*,*KMT2C*, *ARID1A*) and oncogenes (*KRAS*, *PIK3CA*, *ERBB2*, *CTNNB1*)(1,8). Particularly intriguing is the emergence of *TP53* and *KRAS* mutations as early events in gallbladder carcinogenesis, suggesting their fundamental role in malignant transformation (9,10). However, the precise mechanisms by which these genetic alterations drive GBC development remain elusive, largely due to the absence of appropriate experimental models that can recapitulate the early stages of disease progression.

To address these critical gaps, we present here a novel approach to modeling GBC through the engineering of a murine cell line, mGBC1-ZH, derived from normal mouse gallbladder organoids expressing clinically relevant genetic drivers. This model enables both subcutaneous and orthotopic tumor growth in immunocompetent hosts, maintains key pathological and molecular features of human GBC. Through comprehensive genomic and transcriptomic characterization, we trace the molecular evolution from normal gallbladder cells to transformed cancer cells, providing valuable insights into the early events in GBC development. Furthermore, we found that epithelial-derived neutrophil-activating peptide-78 (ENA78/CXCL5) was elevated in GBC and investigated its biological functions in proliferation, metastasis, and neutrophil recruitment using the syngeneic mouse model. We also demonstrated the utility of this model for evaluating therapeutic responses in an intact immune environment, establishing a valuable platform for both mechanistic studies and therapeutic development.

## Materials and Methods

### Sex as a biological variable

Male and female mice were used for all experiments. Data show similar findings for both sexes.

### Murine gallbladder organoid culture

Gallbladders were harvested from *Kras^LSL-G12D/+^/Trp53^fl/fl^* male mice and washed with pre-chilled PBS. Tissues were minced into approximately 0.5 ∼ 1 mm^3^ pieces using fine scissors. The minced tissue was transferred into a 15 ml centrifuge tube and washed with ice-cold DMEM/F12 medium (OPM Biosciences, China) containing 1 % BSA (Sigma, USA). Tissue fragments were collected by centrifugation at 200 g for 5 min at room temperature and dissociated in 5 ml of mixed dispase (Suzhou Jiyan Biotech. Co. Ltd., China) containing 10 μM Y27632 (Selleck, USA) with shaking at 37°C for 15 ∼ 20 min. The cell suspension was centrifuged at 300 g for 5 min at 4 °C and the pellet was washed with 10 ml cold DMEM/F12 medium containing 1 % BSA. After removing the supernatant, the cell pellet was resuspended with 100 μl murine gallbladder organoid medium (MOCM009, Suzhou Jiyan Biotech. Co. Ltd., China) containing 10 μM Y27632. The cell suspension was mixed with 100 μL cold growth factor-reduced Matrigel (Corning, USA) on ice. Fifty μl of the mixture containing 2,000 cells was seeded in the center of each well of a low-adherent 24-well plate and incubated at 37°C with 5% CO2 for approximately 20 min until Matrigel solidification. Each droplet was then overlaid with 500 μL mGOM containing 10 μM Y27632 for 24 h, followed by replacement with fresh mGOM medium without Y27632. Medium was changed every 2-3 days. Organoids were cultured for 7-10 days after initial isolation and subsequently passaged every 5-7 days by mechanical dissociation using a P200 pipette tip.

### Lentiviral transduction of gallbladder organoids

Established gallbladder organoids were dissociated to single cells using TrypLE express solution (Thermo Fisher, USA). A cell suspension containing 1×10^5^ cells in 200 μl medium was incubated with VSVG-LENTAI-UbiC-Cre-EGFP-WPRE-pA (LTR) (Taitool Bioscience, Shanghai, China) following the manufacturer’s protocol at 37°C with 5% CO2 for 12 h. Transduced cells were replated following the organoid culture protocol described above. Successfully transduced organoids reformed in Matrigel and showed EGFP expression within 48 h post-transduction. Representative images were captured using a fluorescence microscope (Leica). Validation of Cre-mediated recombination was performed by PCR following published protocols (11,12). All primer sequences are listed in Supplementary Table S4.

### Establishment of mGBC1-ZH cell line

After 5 passages, transduced gallbladder organoids were dissociated to single cells and plated in 12-well plates for two-dimensional (2D) culture. The culture medium consisted of a 1:1 mixture of mGOM and DMEM/F12 supplemented with 10% Fetal Bovine Serum (FBS, Gibco, USA). After three passages, cells were transitioned to DMEM supplemented with 10% FBS for continued culture. Following 15 passages in 2D culture, cells were injected subcutaneously into mice. Resultant subcutaneous tumor was harvested at two weeks post-injection and dissociated using 1 mg/ml type I collagenase, 1 mg/ml type IV collagenase, and 0.5 mg/ml DNase I. Isolated cells were subsequently cultured in DMEM/F12 supplemented with 10% FBS for 20 generations. Cultures were tested for mycoplasma contamination weekly.

### Immunohistochemistry and immunofluorescence

Gallbladder organoids were fixed in 4% paraformaldehyde (PFA) for 4 h at room temperature (RT) following Matrigel removal. Organoids were then embedded in a special embedding gel (SE-gel, Suzhou Jiyan Biotech. Co. Ltd., China) and incubated for 30 min at 37°C until solidification. The SE-gel-embedded organoids were subsequently fixed in 4% PFA overnight at 4°C before paraffin embedding. Tumor tissues were similarly fixed in 4% PFA and embedded in paraffin. All embedded samples were sectioned at 5 μm thickness and processed for hematoxylin and eosin (H&E) staining and immunohistochemistry (IHC) following standard protocols. For IHC, the following primary antibodies were used: anti-CK7 (GB115696, Servicebio, 1:250), anti-CK19 (GB11197, Servicebio, 1:100), anti-Ki67 (GB111141, Servicebio, 1:250), anti-CD3 (GB12014, Servicebio, 1:500), anti-CD20 (GB11540, Servicebio, 1:800), anti-CD31 (GB113151, Servicebio, 1:300), anti-Ly6G (GB11229, Servicebio, 1:400), and anti-MPO (GB12224, Servicebio, 1:1000). Images were acquired using a digital slide scanner (Pannoramic 250 FLASH III, 3DHISTECH Ltd).

Immunofluorescence (IF) staining was performed as followed. Cells were seeded on coverslips and cultured for 24 h, then fixed in 4% PFA for 10 min at room temperature. Following fixation, cells were permeabilized with 0.5% Triton X-100 for 10 minutes and blocked with 3% BSA for 30min. Primary antibodies were applied overnight at 4°C in a humidified chamber, followed by secondary fluorescence-conjugated antibodies for 1 hour at room temperature. Nuclei were stained with DAPI for 5 minutes. In addition to the primary antibodies used for IHC, anti-EpCAM (GB11274, Servicebio, 1:100) was also used for IF staining. Images were acquired using a digital slide scanner (Pannoramic MIDI, 3DHISTECH Ltd).

### Development of subcutaneous and orthotopic tumors

Six to eight-week-old female C57BL/6J mice were obtained from Gempharmatech (Shanghai, China). mGBC1-ZH cells were dissociated using 0.25% TrypLE, and a single-cell suspension containing 2×10^6^ cells in 0.1 ml was injected subcutaneously into the right inguen of each mouse. Tumor growth was monitored starting 7 days post-injection. Tumor size was measured every three days with a caliper, and tumor volume was calculated using the formula: (shortest diameter)^2^ × longest diameter × 0.5.

Orthotopic implantation was performed following the method described by Smith et al(13). with minor modifications. Mice were fasted for 4 hours and anesthetized with chloraldurat. For implantation, approximately 2×10^5^ cells in 10 μl DPBS with 10 μl Matrigel were used for each mouse. The gallbladder was exposed through a 0.8-1.0 cm abdominal midline incision and punctured with a 31G insulin syringe. Following bile extrusion and cleaning, 20 μl of cell suspension was injected into the gallbladder using a 29G insulin syringe. The syringe was removed after solidification of the suspension. Following suturing of abdominal wall, mice were placed on a 37°C heating pad for recovery. Mice were monitored daily for the first week, and subsequently every three days for three weeks. At four weeks post-implantation, mice were euthanized by CO_2_ exposure, and primary tumors were excised (Supplementary Figure 2C). The study was approved by the Ethics Committee of Fudan University School of Basic Medical Sciences and a maximal tumor diameter < 20 mm was permitted. We monitored that the maximal tumor diameter was not exceeded.

### Karyotyping analysis

Chromosomal analysis was performed on established tumor cells during exponential growth phase. Cells were treated with 0.1 μg/ml colcemid for 1.5 h and processed using standard air-dried methods. Cytogenetic analysis was conducted on G-banded metaphase cells, with a total of 20 cells examined.

### Short tandem repeat (STR) analysis

Genomic authentication was performed by amplifying eighteen short tandem repeat (STR) loci using multiplex PCR. The human-specific marker (Human TH01) was included to screen for potential human cell contamination. Samples were analyzed using an ABI Prism® 3130 XL Genetic Analyzer, and data were processed with Gene Mapper® ID v3.2 software (Applied Biosystems). Appropriate positive and negative controls were run and validated.

### Human tissue microarray analysis

To assess the prognostic effect of CXCL5, a paraffin GBC tissue microarray was acquired from Zhongshan Hospital (Fudan University, Shanghai, China)with obtainedinformed consent from all patients(14). IHC staining was performed as described above.QuPath (v0.5.1) was used to analyzed the IHC intenstiy.Overall survival (OS) were calculated by the Kaplan-Meier method and differences were analyzed by the log-rank test.

### Human GBC Cell culture

The OCUG-1 cell linewere cultured in DMEM medium containing 10% fetal bovine serumand 1% penicillin/streptomycinat 37 □ with5% CO2 in an incubator.Mycoplasma testing was performed each week.

### Lentivirus infection

Cells were infected with lentivirus to deliver shRNA-control (shNT), shRNA-CXCL5 (shCXCL5) constructs, using lentiviral preparations purchased from Genechem(Shanghai, China). Plasmids for CXCL5 overexpression were sourced from the same supplier. Virally infected cells were selected by5mg/ml puromycin for 96 hours.

### Real-time quantitative PCR

Total RNA was isolated from cells with TRIzol Reagent (Invitrogen) following the manufacturer’s protocol. First-strand cDNA was synthesized from the extracted RNA using the PrimeScript^TM^ RT Reagent Kit with gDNA Eraser (Takara). Quantitative real-time PCR (qPCR) was subsequently performed on a Roche LightCycler 480 Real-Time PCR System using ChamQ SYBR qPCR Master Mix (Vazyme). Gene expression levels were relatively quantified using the 2^^-ΔΔCt^ method. The primers used for qPCR analyses are as follows:

Human CXCL5 Forward: 5’-GACGGTGGAAACAAGGAAAA-3’, Reverse: 5’-GCTTAAGCGGCAAACATAGG -3’

Mouse Cxcl5 Forward: 5’-CTGGCATTTCTGTTGCTGTTCA-3’,Reverse: 5’-GGGATCACCTCCAAATTAGCGA -3’

Human GAPDH Forward: 5’-CCATGTTCGTCATGGGTGTGA-3’, Reverse: CATGGACTGTGGTCATGAGT

Mouse Gapdh Forward: 5’-AGGTCGGTGTGAACGGATTTG-3’, Reverse: TGTAGACCATGTAGTTGAGGTCA

Human CD66b Forward:5’-GCGAGTGCAAACTTCAGTGACC-3’, Reverse: ACTGTGAGGGTGGATTAGAGGC

### Cell proliferation and colony formation assays

For proliferation assays, established tumor cells were seeded in 96-well plates at either 2,000 cells per well for growth rate assessment or at 10,000 cells per well for drug treatment studies, in 100 µl culture medium. Cell viability was measured using a Cell Counting Kit-8 (Beyotime Biotechnology) assay either 72 h after chemical reagent treatment or every 24 h after plating. Following 1□h incubation at 37□, absorbance was measured at 450□nm using a microplate reader (BioTek, Vermont, USA). All experiments were performed in triplicate.

### Migration and invasion assays

For wound-healing assays, mGBC1-ZH cells were seeded in 24-well plates (2 × 10^5^ cells/ml) and cultured until confluence. Wounds were created using a 1 ml micropipette tip, and cells were washed with PBS before addition of fresh growth medium containing FBS. Wound closure was monitored at 0, 24, and 48 hours post-wounding and quantified using ImageJ software. Wound closure percentage was calculated as [(initial area - final area)/initial area] × 100.

Invasion assays were performed using transwell chambers (Corning, USA). Cell suspensions (3 × 10^4^ in 200 μl) were seeded in the upper chamber, while 600 μl of fresh growth medium containing 10% FBS was added to the lower chamber. After 24 h incubation at 37 °C with 5% CO_2_, invaded cells were stained with Giemsa (Sigma, USA) and imaged using a bright-field microscope.

### RNA-seq data analysis

RNA-seq reads were trimmed and pseudoaligned to the mouse transcriptome (GENCODE v35, GRCm38) using kallisto (v0.46.1)(15) to quantify gene expression. Differential expression analysis of protein-coding genes were performed using DESeq2 (v1.28.1), comparing samples pair wisely or using a linear regression model. Differentially expressed genes (DEGs) were identified using the thresholds of fold change >2 or <0.5 and an FDR < 0.05. Gene Set Enrichment Analysis (GSEA) was performed using clusterProfiler (v4.2.2) to evaluate both GO and KEGG pathways, with an adjusted P-value < 0.05 considered significant. Temporal gene expression patterns were clustered and annotated using ClusterGVis (https://github.com/junjunlab/ClusterGVis).

### Analysis of Whole Exome sequence data

We aligned paired-end whole-exome sequencing reads to the mouse reference genome (GRCm38) using BWA memwith default parameters, and generated BAM files with samtools. We then optimized the alignments according to GATK (v4.1.0.0) best practices. We used MarkDuplicates to identify duplicate artifacts, and BaseRecalibrator with knownSites set to MGP.v5.snp_and_indels.exclude_wild.chr.vcf.gz (Sanger Mouse Genomes Project; https://ftp.ebi.ac.uk/pub/databases/mousegenomes/REL-1505-SNPs_Indels/) to account for systemic base quality errors. For variant calling, we used MuTect2under the tumor-only mode. We filtered the variants using FilterMutectCalls with --max-events-in-region set to 3. Then we filtered the mutations with the following criteria: (i) “PASS” in the Filter label, (ii) the variant allele frequency greater than 0.05, and (iii) the reads supporting the locus greater than 5.

To determine copy number variations, we used CNVkit (v0.9.10) with the tumor-only options, using the batch option with the following parameters: cnvkit.py batch [recalibrated and sorted bam file] -normal --targets [exome regions bed file] --fasta [GRCm38] --annotate [http://hgdownload.soe.ucsc.edu/goldenPath/mm10/database/refFlat.txt.gz] --output-reference [SAMPLE].cnn.

### Hierarchical clustering

Transcriptome datasets of 20 human biliary tract cancers cell lines were obtained from previously report (16). mRNA abundance expressed in transcripts per million (TPM) was compared for mGBC1-ZH and human biliary tract cancers cell lines. Unsupervised hierarchical clustering was generated using Morpheus (https://software.broadinstitute.org/morpheus). Heatmap was generated using the 400 significant differential expression genes between the two subtypes.

### Correlation analysis of DEGs in human and mouse GBC

We identified DEGs between malignant and non-malignant epithelial cells using single-cell RNA sequencing data of human GBC samples(17). The fold change values of 503 genes that were consistently dysregulated in both human and mouse RNA-seq datasets were obtained. Pearson’s correlation analysis was performed to assess their cross-species transcriptional similarity.

### In vivo drug response studies

mGBC1-ZH cells (2×10^6^ cells) were injected subcutaneously into the right inguen of each mouse. After 10 days, when tumors were established, mice (n = 20) were randomly assigned into four treatment groups with equivalent mean tumor volumes. Each group received the following regimens: control group (200μl of 0.9% saline, twice weekly), gemcitabine group (30 mg/kg, twice weekly), and cisplatin group (7.5 mg/kg cisplatin, once weekly). All treatments were administered intraperitoneally. After three cycles of treatment, mice were euthanized by CO_2_ exposure and tumors were harvested. Tumor volumes and weights were measured, and data are presented as the means ± S.D.

## Results

### In vitro generation of gallbladder organoids with Kras activation and Trp53 deletion

To model human GBC tumorigenesis, we generated mouse gallbladder organoids with concurrent *Kras* activation and *Trp53* (encoding mouse *p53*) deletion (Figure 1A). First, we established organoid cultures from gallbladder epithelium isolated from *Kras^LSL-G12D/+^/Trp53^fl/fl^*; C57BL/6J male mice. These organoids exhibited thin-walled cystic structures (Figure 1B). Histological analyses revealed that the organoids maintained a monolayer epithelial cystic architecture, closely resembling the original tissue architecture (Figure 1B). The organoids strongly expressed biliary epithelial markers CK7 and CK19, and multiple proliferating epithelial cells as indicated by Ki67 immunostaining (Figure 1B). Subsequently, we introduced a lentiviral vector expressing Cre recombinase and EGFP into the organoids to activate *Kras*^G12D^expression and delete *Trp53* gene (Supplementary Figure 1A). Successful genetic recombination was confirmed by conditional PCR (Supplementary Figure 1B). The resulting *Kras*^G12D^/*Trp53*^-/-^ organoids developed irregular cystic structures with thicker walls compared to normal organoids (Figure 1B). These genetically modified organoids maintained expression of CK7 and CK19, while displaying enhanced Ki-67 staining relative to normal organoids, indicating increased proliferative capacity (Figure 1B).

**Figure. 1.**
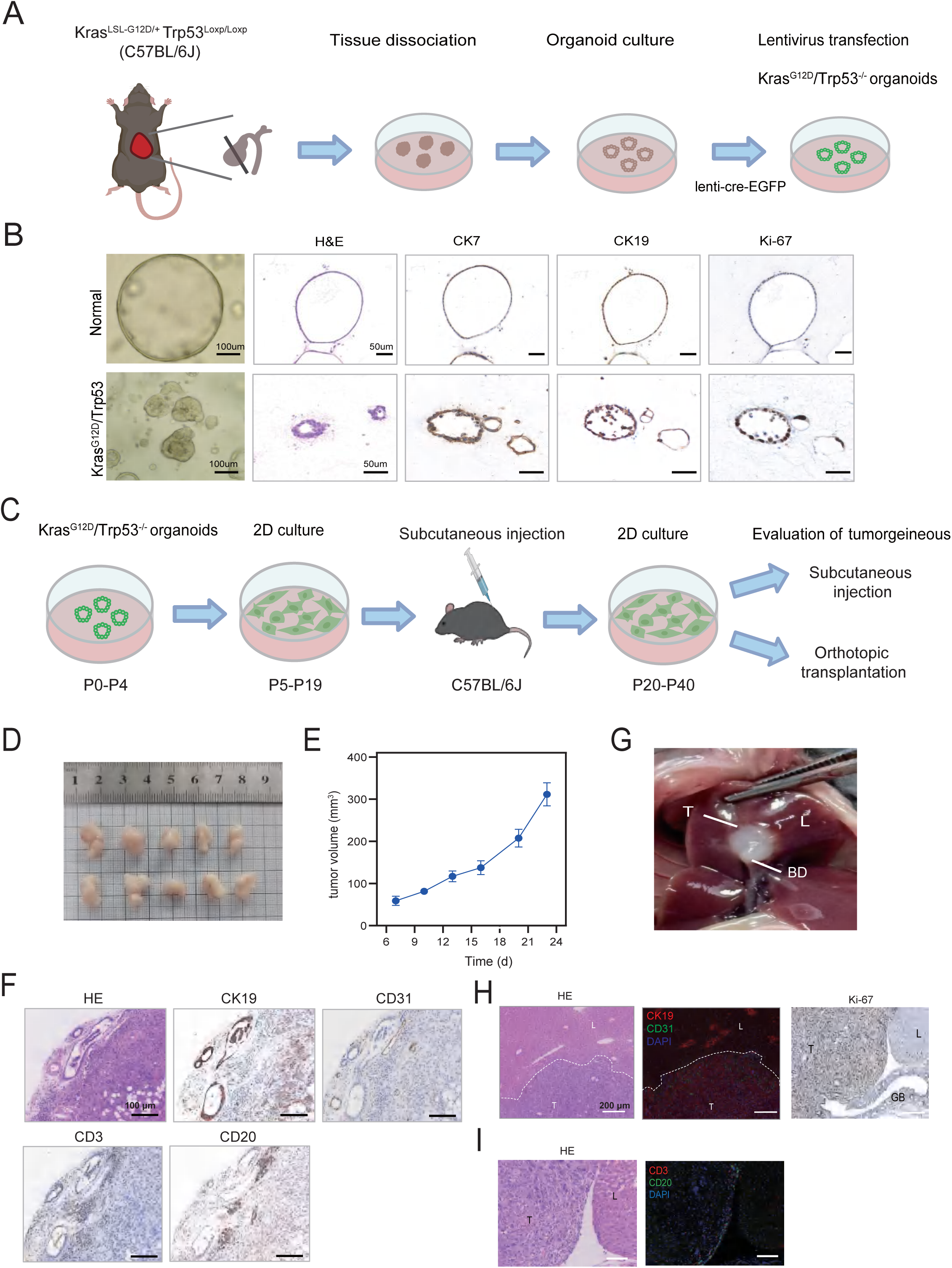
Generation of murine GBC model in syngeneic wild-type mice. **(A)** Schematic of the strategy for generating *Kras*^G12D^/*Trp*53^-/-^ organoids. Gallbladder of C57BL/6J mouse was isolated and processed as described in the Methods. **(B)** Representative bright field, H&E and immunohistochemical staining for CK7, CK19 and Ki-67 of the normal organoids and the *Kras*^G12D^/*Trp53*^-/-^ organoids. Scale bars: 100 μm (bright-field image) and 50 μm (H&E and immunohistochemical staining). **(C)** Schematic of the strategy for generating gene edited GBC cell line and syngeneic mice model. **(D-E)** The subcutaneous tumors **(D)** and tumor growth curve **(E)** monitored (n = 10). Data were presented as mean ±SD. **(F)** Representative H&E, immunohistochemical for CK19, CD31, CD3 and CD20 of subcutaneous tumor. Scale bars: 100 μm. **(G)** Macroscopic view of an orthotopic gallbladder tumor. T, tumor; L, liver; BD, bile duct. **(H)** Representative H&E, Immunofluorescence staining for CK19, CD31 of orthotopic tumor. Direct invasion of cancer cells into the liver tissue (white dotted line). Scale bar, 200 μm. **(I)** H&E, and immunofluorescence for CD3 and CD20 of orthotopic tumor.

### Establishment of a murine GBC model in syngeneic wild-type mice

After successfully engineered *Kras*^G12D^/*Trp53*^-/-^ organoids, we next sought to develop a murine GBC cell line that could be used for long-term studies. This would address a significant gap in the field, as no widely used murine GBC cell lines have been reported in the literature.The modified organoids were initially cultured in a three-dimensional (3D) environment for four passages to ensure their growth and stability. Following this, they were transferred to a two-dimensional (2D) culture system where they were maintained for an additional 15 generations (Figure 1C). The initial epithelial cells exhibited a polygonal and tightly connected morphology with many multinuclear cells (Supplementary Figure 2A).

To assess their tumorigenic potential, we injected these cultured cells into the subcutaneous tissue of syngeneic wild-type (WT) mice. Among three mice tested, only one developed a palpable tumor after two weeks. To enhance the tumorigenicity of the cells, the resulting subcutaneous tumor was isolated, digested, and cultured in vitro for an additional 20 generations (Figure 1C). During this process, these cells underwent significant morphological changes, becoming elongated and spindle-shaped, suggesting an enhanced epithelial-mesenchymal transition (EMT)(Supplementary Figure 2A). Immunofluorescence analysis demonstrated expression of CK7, CK19, and EpCAM, confirming their epithelial origin. Strong Ki-67 expression was observed, indicating a high proliferation rate (Supplementary Figure 2B). We designated this newly established cell line as mGBC1-ZH.

To evaluate the tumorigenic capacity of mGBC1-ZH cells under an intact immune system, we performed subcutaneous injections into syngeneic WT mice. These cells formed palpable tumors within one week in all 10 mice tested (Figure 1D), demonstrating their robust tumorigenic potential. Longitudinal monitoring of tumor growth revealed a rapid increase in size (Figure 1E), indicating aggressive proliferation in vivo. Histological analysis of the subcutaneous tumor revealed consistent expression of CK19(Figure 1F). Importantly, we observed abundant CD31-positive vascular structures throughout the tumor, indicating extensive angiogenesis that likely supports the high metabolic demands of these rapidly proliferating cells(Figure 1F). Analysis of the tumor immune microenvironment revealed aggregation of CD3+ T cells and CD20+ B cells in the tumor margins (Figure 1F), characteristic of tertiary lymphoid structures. Notably, these immune cells were sparsely distributed within the tumor core, indicating a “cold” tumor microenvironment, a feature frequently observed in human GBC. These findings demonstrate that mGBC1-ZH cells can establish tumors that recapitulate key features of human GBC in immunocompetent hosts.

To further characterize this model, we assessed the orthotopic tumorigenic capacity of mGBC1-ZH cells. We injected cells resuspended in matrigel into the gallbladder lumen of syngeneic WT mice (Supplementary Figure 2C). After four weeks, the animals were sacrificed, revealing solid orthotopic tumors (Figure 1G). Histopathological analysis revealed amoderately differentiated adenocarcinoma with liver invasion (Figure 1H). Consistent with the subcutaneous tumors, the orthotopic tumors maintained CK19 expression and exhibited higher Ki-67 levels than normal gallbladder tissue (Figure 1H). Similar to human GBC, neo-angiogenesis and lymphocyte infiltration were observed primarily at the tumor-liver interface (Figure 1H, 1I). Notably, in one orthotopic tumor, we observed a distinct cellular architecture in which tumor cells were encapsulated by stromal cells, which appeared to impede immune cell infiltration into the tumor core (Supplementary Figure 2D). This structure closely resembled the tumor immune barrier (TIB) previously described in hepatocellular carcinoma (18), suggesting potential shared immune evasion mechanisms between GBC and other gastrointestinal cancers.

### Genetic alterations of mGBC1-ZH cell line

After confirming of its in vivo tumorigenic capacity, we performed comprehensive characterization of the mGBC1-ZH cell line. Short Tandem Repeat (STR) profiling confirmed the cell line’s authenticity and origin (Figure 2A). Karyotyping analysis revealed significant chromosomal instability, characterized by both numerical and structural abnormalities (Figure 2B). Chromosomal analysis revealed counts ranging from 53 to 80, with most cells containing 60 to 70 chromosomes. This aneuploidy, a hallmark of cancer cells, is frequently observed in *TP53*-mutant tumors that display similar copy number alterations (19,20).

**Figure. 2.**
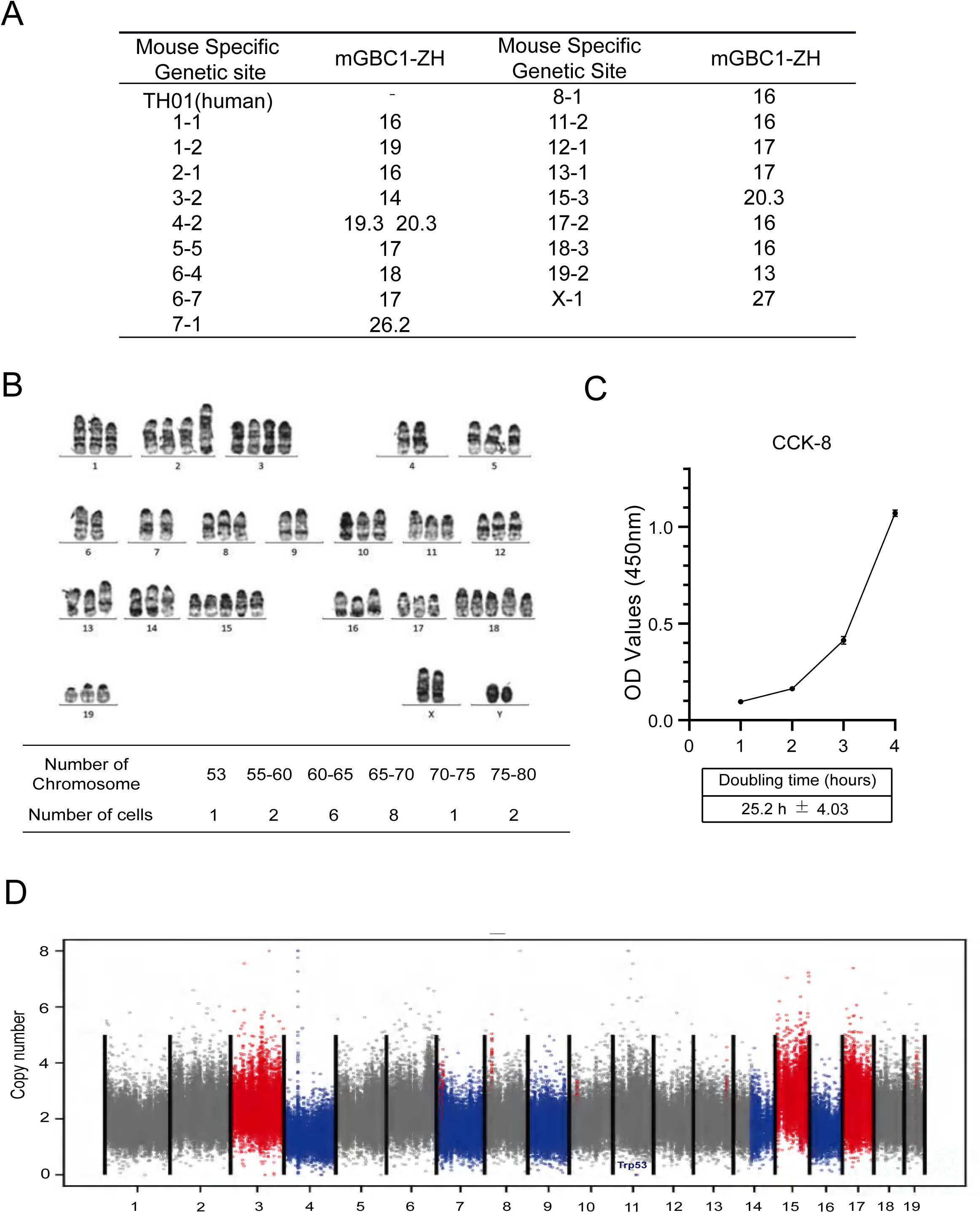
Genetic characteristics of mGBC1-ZH cell line. **(A)** Short tandem repeat (STR) DNA profling of mGBC1-ZH. **(B)** The distribution of chrmosome number analyzed by karotyping of mGBC1-ZH (n=20). A representative karyotype of a metaphase spread prepared from mGBC1-ZH was shown. Karyotype description: 63, XXYY, +2, +3, -4, -6, -7, -9, +15, +15, +18, +18, t(2;14) (G;C3). **(C)** Growth curve of mGBC1-ZH cells at 24, 48, 72 and 96h after plating. Data were presented as mean ±SD. **(D)** Copy number alterations in mGBC1-ZH. Grey, red and blue dots represent base line, copy number gain and loss, respectively.

mGBC1-ZH cells exhibited robust proliferation with an average doubling time of 25.2 ± 4.03 h (Figure 1C). To characterize the genetic landscape of mGBC1-ZH, we performed whole-exome sequencing (WES) on the established cell line (P40) and matched normal gallbladder organoids. The analysis revealed 193 somatic mutations and variants, with missense mutations (n=179) constituting the vast majority (Supplementary Figure 3A, Supplementary Table S1). Single-nucleotide variants represented the predominant variant type (Supplementary Figure 3B). Mutational signature analysis demonstrated a characteristic pattern of G > A/C > T and T > C/A > G transversions (Supplementary Figure 3C), consistent with mutational signatures previously documented in human GBC specimens(7,8). Copy number analysis identified recurrent chromosomal alterations, including losses on chromosomes 4, 7, 9, and 16, and concurrent gains on chromosomes 3, 15, and 17 (Figure 2D). Notably, these alterations encompassed loci containing established GBC-driver genes (6), including deletion of the *Cdkn2a* locus (chromosome 4) and amplification of *Pik3ca* (chromosome 3) and *Vegfa* (chromosome 17) loci. We also detected deletion of the *Trp53* locus on chromosome 11, consistent with the engineered *Trp53* mutation. This pattern of chromosomal aberrations parallels those documented in murine models of pancreatic ductal adenocarcinoma (PDAC) (19), where sporadic p53 loss of heterozygosity preceded malignant transformation and was associated with similar chromosomal instability. This observation suggests a conserved mechanism of genomic evolution driven by p53 inactivation across distinct gastrointestinal malignancies.

### Transcriptional evolution in syngeneic murine GBC model generation

To understand the molecular evolution during syngeneic murine GBC model development, we performed transcriptomic profiling at the following key stages: initial normal gallbladder organoids, early-passage cells following 2D adaptation (P7), and established GBC cells (P40). Principal component analysis (PCA) demonstrated clear transcriptional segregation of the established P40 cells from both normal organoids and P7 cells (Supplementary Figure 3D). The maintenance of gallbladder lineage identity throughout model development was confirmed by robust expression of gallbladder and biliary tract markers, including *Aldoa*, *Krt19* (*CK19*), *Krt7* (*CK7*), *Epcam*, and *S100a6*. Importantly, these cells maintained tissue specificity, showing minimal expression of hepatocellular carcinoma markers (*Afp* and *Gpc3*) and hepatocyte markers (*Alb*, *Ttr*, *Apoa1*, and *Apoe*), as illustrated in Figure 3A.

**Figure. 3.**
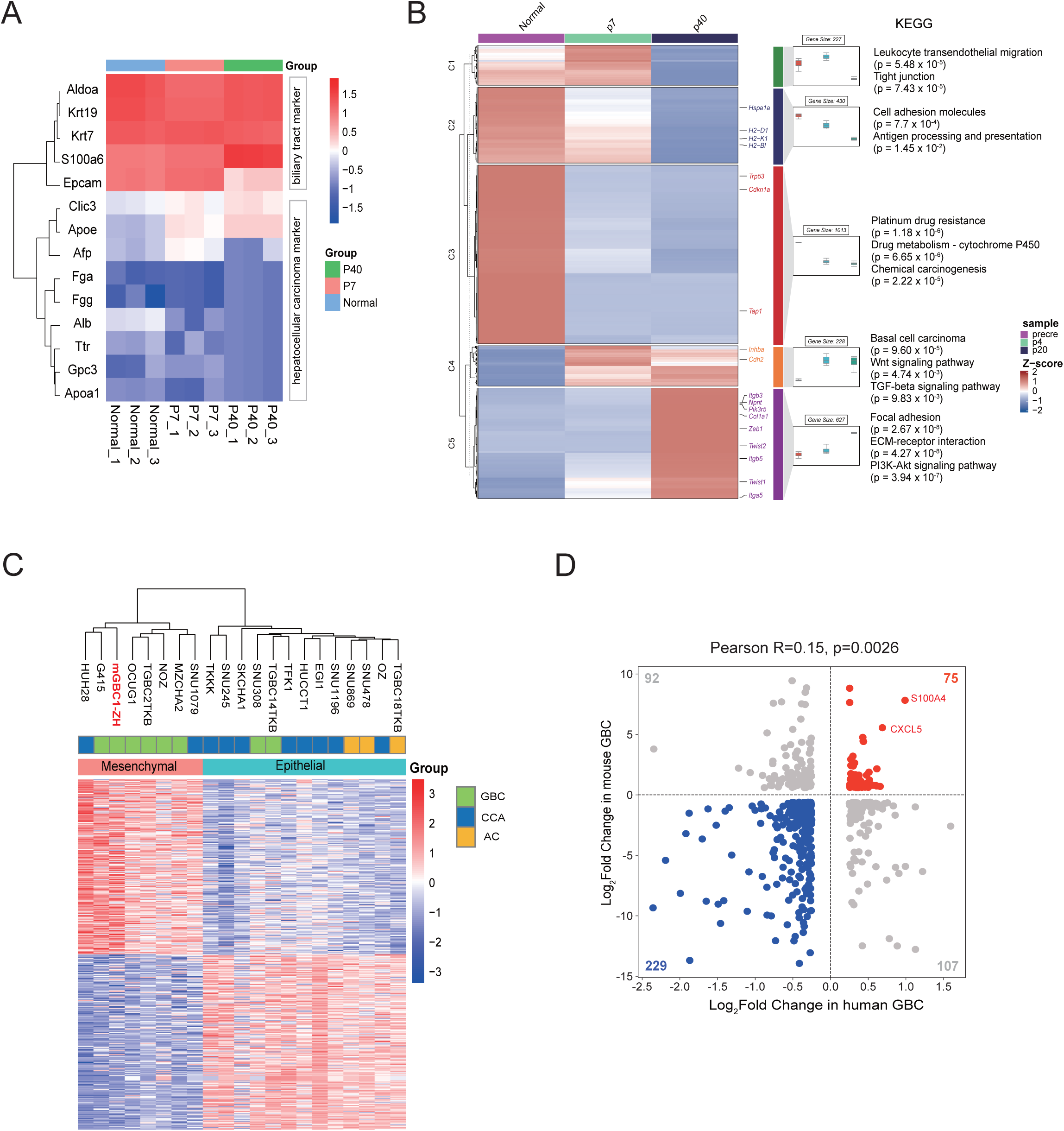
Gene exprssion profiles of the mGBC1-ZH cell line. **(A)** Heatmap showing the gene expression profiles associated with biliary tractand hepatocellular carcinoma in normal organoids, P7 and P40 cells. **(B)** Heatmap representation of time course RNA-seq data showed rapid activation of pathways typical of Wnt and TGF-beta and down-regulation of pathways characteristic of platinum drug resistance and p53 signaling. **(C)** Unsupervised hierarchical clustering of mGBC1-ZH cells and human biliary tract cancers cell lines based on basal level expression of all genes. Heatmap shows the 400 genes with significant differential expression between the two subtypes. GBC, gallbladder carcinoma; CCA, cholangiocarcinoma; AC, ampullary cancer. **(D)** Correlation analysis of DEGs (503 genes) identified in both human and mouse GBC. Pearson’s correlation analysis was used to analyze the correlation.

Comparison of transcriptional profiles revealed 4,872 differentially expressed genes (DEGs) between P7 cells and normal organoids (Supplementary Table S2), with 2,367 upregulated and 2,505 downregulated genes. Analysis of the P40 cells and normal organoidsidentified 5,588 DEGs (Supplementary Table S3), including 2,312 upregulated and 3,276 downregulated genes. Importantly, P7 and P40 cells shared extensive transcriptomic similarities, with 1,195 commonly upregulated genes and 1,696 commonly downregulated genes, suggesting conserved transcriptional changes during model establishment(Supplementary Figure 3E). Pathway analysis revealed significant upregulation of the PI3K-Akt and MAPK signaling pathways in both P7 (Supplementary Figure 3F)and P40 cells (Supplementary Figure 3G), consistent with the characteristic signaling patterns of KRAS-mutated cancers(21). Additionally, both P7 and P40 cells showed upregulation of ECM-receptor interaction and focal adhesion pathways, which regulate cell-matrix interactions, migration, and invasion(22). In contrast, genes involved in gallbladder-specific functions, including glutathione metabolism and metabolism of xenobiotics by cytochrome P450, were downregulated. The p53 signaling pathway was significantly downregulated in P40 cells, aligning with the introduced *Trp53* mutation. These findings demonstrate progressive transcriptional reprogramming during syngeneic GBC model generation, with both early and late passages showing distinct molecular signatures compared with normal organoids.

To elucidate the temporal transcriptional changes during model development, we classified all DEGs between time points into five distinct patterns (Cluster 1-5) using hierarchical clustering (Figure 3B). Genes involved in focal adhesion, ECM-receptor interaction, and PI3K-Akt signaling pathways showed a gradual increase in expression from P7 to P40 (cluster 5), whereas genes associated with cell adhesion molecules and antigen processing and presentation pathways exhibited continuous downregulation (cluster 2). Notably, genes involved in Wnt and TGF-β signaling pathways displayed rapid upregulation at the early stages (cluster 4), suggesting their potential roles as critical drivers of tumor initiation and early progression. Genes associated with platinum drug resistance and p53 signaling showed early downregulation at P7 (cluster 3), which was consistent with the *Trp53* mutation. These distinct temporal expression patterns provide insights into the molecular events during the progressive transformation of normal gallbladder organoids into syngeneic GBC model.

### Similar transcriptome between mGBC1-ZH and human GBC

To assess whether the mGBC1-ZH murine model recapitulates molecular features of human GBC, we performed comparative transcriptomic analyses with established human systems. In a recent study, David*et al.* analyzed the transcriptomic profile of 20 human biliary tract cancers cell lines, resulting in their classification into two major subtypes that differ in epithelial and mesenchymal characteristics (16). Unsupervised hierarchical clustering positioned mGBC1-ZH (P40) within the mesenchymal subgroup (Figure 3C), a classification corroborated by significant enrichment of EMT hallmark gene signatures. Notably, this mesenchymal cluster demonstrated gallbladder-specific predominance, with 5 out of 7 cell lines originating from human GBC specimens, suggesting conserved transcriptomic features between murine and human GBC cell lines.

To further validate this cross-species similarity, we analyzed single-cell RNA sequencing data from human gallbladder carcinomas (17). By comparing 503 DEGs identified in both human malignant epithelial cells (vs. normal) and mGBC1-ZH cells (vs. murine gallbladder controls), we observed significant transcriptional concordance (Pearson correlation coefficient r = 0.15, *p* = 0.0026; Figure 3D),confirming their similarity at the transcriptomelevel.

### CXCL5 is upregulated in human GBC and promotes cell proliferation,migration, and invasion

We noticed that *CXCL5* was elevated in both mGBC1-ZH and human malignant epithelial cells, and was upregulated as early as P7 cells (Supplementary Figure 4A). To further investigate the potential role of CXCL5 in GBC, we first examined the expression of *CXCL5* in paired GBC tumor tissues and adjacent non-tumor tissues. RT-qPCR showedan elevated expression of CXCL5in GBC tumor tissues compared with the normal tissues(Figure 4A),thiswas further confirmed by immunohistochemistry staining(Figure 4B).Next, to determine the association of CXCL5 with the prognosis of patients in GBC, we performed immunohistochemistry on the tissue microarrays established from 167 GBC tissues. The result showed that higher CXCL5 intensity in tumors associated with worse prognosis (*p*=0.014, Figure 4C).

**Figure. 4.**
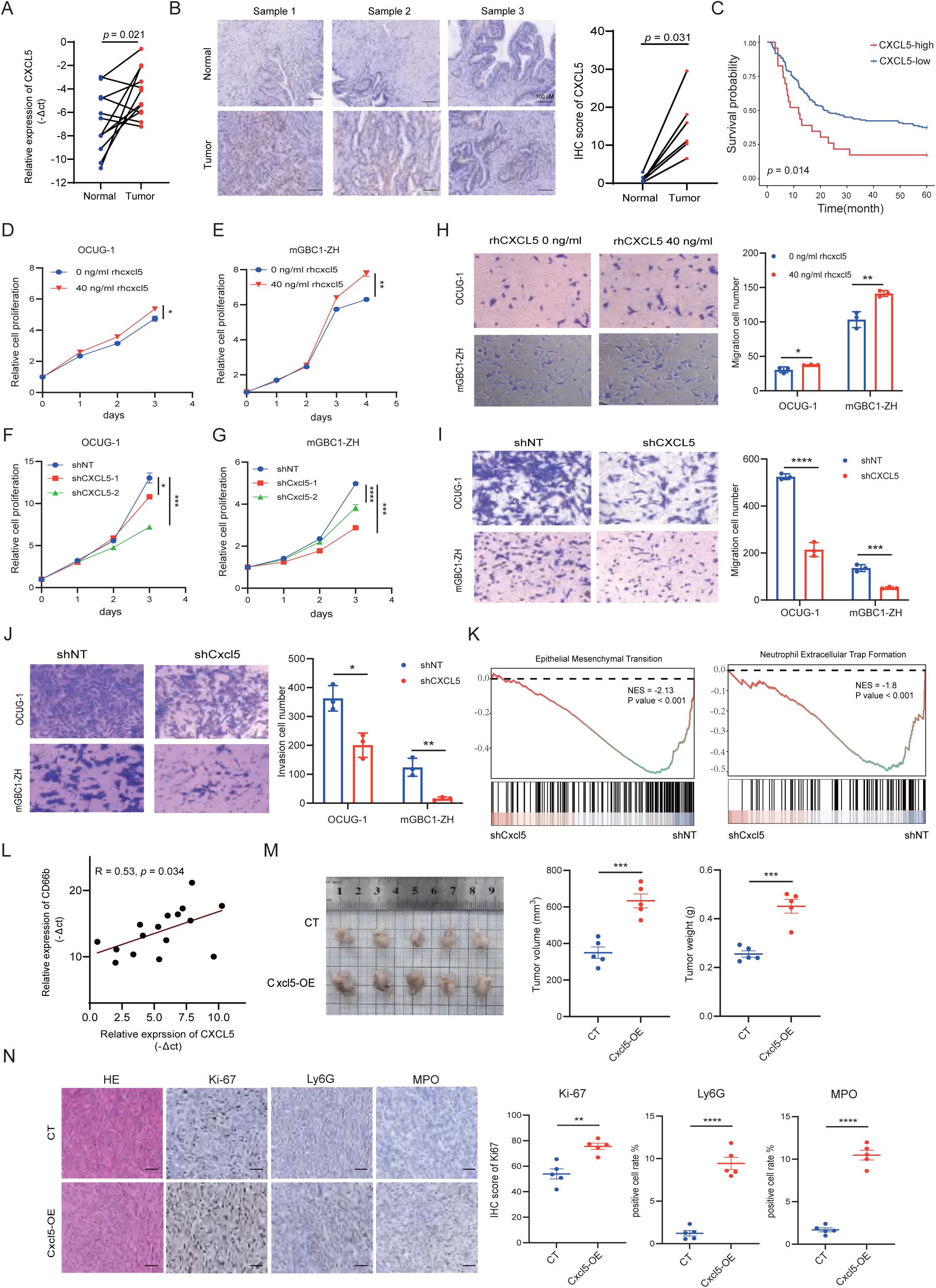
CXCL5 is upregulated in human GBC and enhanced cell proliferation, metastasis and neutrophil recruitment. **(A)** qRT-PCR analyses of CXCL5 expression in gallbladder cancer tissues and paired normal tissues. **(B)** Representative immunohistochemical staining of CXCL5 between GBC tumors and paired normal tissues (scale bars, 100 μm). **(C)** High expression of CXCL5 is associated with poorer overall survival. **(D-E)** Relative proliferative rates of OCUG-1 **(D)** and mGBC1-ZH **(E)** under the treatment of rhCXCL5 or not detected by CCK8 assay. **(F-G)** Relative proliferative rates of OCUG-1 **(F)** and mGBC1-ZH **(G)** cells upon CXCL5 knock down. **(H-I)** Transwell migration assays for OCUG-1 and mGBC1-ZH upon rhCXCL5 treatment **(H)** and CXCL5 knock down **(I)**. **(J)** Transwell invasion assays for OCUG-1 and mGBC1-ZH upon CXCL5 knock down. **(K)** GSEA results based on the RNA-Seq data. **(L)** qPCR analysis of the correlation of CD66b and CXCL5 in human GBC tumors. **(M)** Representative images and quantification of subcutaneous mGBC1-ZH tumors established in C57BL/6J mice. **(N)** Representative images and quantification of H&E, immunohistochemical for Ki-67, Ly6G, and MPO of subcutaneous tumors. Scale bars: 100 μm.

Although CXCL5 has been reported to promote tumor cell proliferation and metastasis in various cancers(23,24), its functional role in GBC tumorigenesis remains unclear. To investigate the contribution of CXCL5 to GBC cell proliferation and disease progression, we performed CCK-8 assays in OCUG-1 and mGBC1-ZH cells. The results demonstrated that treatment with rhCXCL5 significantly enhanced cell proliferation compared to the control group (Figure 4D, E). We then established stable CXCL5-knockdown cell lines using shRNA, with knockdown efficiency confirmed by qRT-PCR and ELISA of the culture supernatants (Supplementary Figure 4B-D). CCK-8 assays revealed that CXCL5 knockdown markedly suppressed cell proliferation (Figure 4F, G). Furthermore, Transwell assays were conducted to assess the effect of CXCL5 on GBC cell migration and invasion. rhCXCL5 treatment significantly promoted cell migration (Figure 4H), whereas CXCL5 knockdown substantially impaired both migratory (Figure 4I) and invasive capacities (Figure 4J). Collectively, these findings indicate that CXCL5 facilitates tumor proliferation, migration, and invasion in GBC.

To further explore the potential mechanisms underlying CXCL5-mediated proliferation and migration in GBC, we performed RNA-Seq on shCxcl5 and shNT mGBC1-ZH cells. We identified 571 significantly upregulated and 583 significantly downregulated genes (Supplementary Figure 4E). Gene Ontology (GO) analysis showed that the downregulated genes were enriched in the biological process of extracellular matrix organization, epithelial cell migration, angiogenesis and neutrophil migration. KEGG showed that the downregulated genes were enriched in cytokine-cytokine receptor interaction, TNF signaling pathway, Cell adhesion molecules, PI3K-Akt signaling pathway, and JAK-STAT signaling pathway (Supplementary Figure 4F). In hepatocellular carcinoma, CXCL5 has been shown to activate the PI3K-Akt signaling pathways to promoted proliferation, migration, and invasion(25). Further, GSEA showed that Epithelial Mesenchymal Transition and Neutrophil Extracellular Trap Formation were significantly negatively enriched in shCxcl5 compared to shNT (Figure 4K). As CXCL5 is an effective neutrophil chemoattractant, these results indicated that CXCL5 may regulates the neutrophils infiltration in GBC.

Through qPCR, we found that CXCL5 expression was significantly positively correlated with CD66b in human GBC tumors (Figure 4L). To investigate the function of CXCL5-mediated neutrophils infiltration in GBC, we established stable Cxcl5 over-expression cell lines (Supplementary Figure 4G, 4H). In vivo analysis in C57BL/6J mouse showed that Cxcl5 over-expression significantly increased tumor volume and tumor weight (Figure 4M). H&E and IHC staining confirmed significantly increased malignancy (Ki-67) and neutrophils infiltration (Ly6G, MPO) in Cxcl5-OE tumors (Figure 4N). Taken together, these results demonstrated CXCL5 modulates GBC proliferation, metastasis and promoted neutrophils infiltration.

### Therapeutic response assessment in the syngeneic GBC model

To evaluate the translational potential of our syngeneic murine GBC model, we assessed its response to standard first-line chemotherapeutic agents employed in human GBC treatment through systematic in vitro and in vivo studies.In vitro dose-response analyses establishedhalf-maximal inhibitory concentrations (IC50) for gemcitabine (GEM) and cisplatin (CIS) of 80.17 nM (Figure 5A) and 23.75 μM(Figure 5B), respectively, demonstrating sensitivity to clinicallyrelevant agents. For in vivo validation, we implemented a subcutaneous xenograft study utilizing a randomized, controlled design (Figure 5C). Twenty mice bearing established tumors (10 days post-implantation) were stratified into four treatment groups (n = 5 per group): vehicle control, GEM, CIS, and their combination (GC), with equivalent baseline tumor volumes across groups (Figure 5D). After three cycles of treatment, all drug treatment groups exhibited significantly reduced tumor volumes (Figure 5E, 5F) and weights (Figure 5G) compared to the vehicle control. Although the GC combination demonstrated a trend towards enhanced antitumor efficacy compared to either monotherapy, this difference did not reach statistical significance. Corroborating these findings, immunofluorescence analysis revealed significant reduction in Ki-67 positive cells among CK19+ epithelial cells in all drug-treated groups (Figure 5H), indicating reduced tumor proliferation.Previous studies have reported increased immune cells infiltration following chemotherapyinmultiplecancer (26,27), consistent with this, we found asignificantincrease ofCD3+ T cellsin all drug treatment groupscompared with the vehicle control (Figure 5I).These results validate our model’s pharmacological responsiveness and underscore its utility as a preclinical platform for evaluating novel therapeutic strategies and elucidating mechanisms of drug resistance in GBC within an immunocompetent microenvironment.

**Figure. 5.**
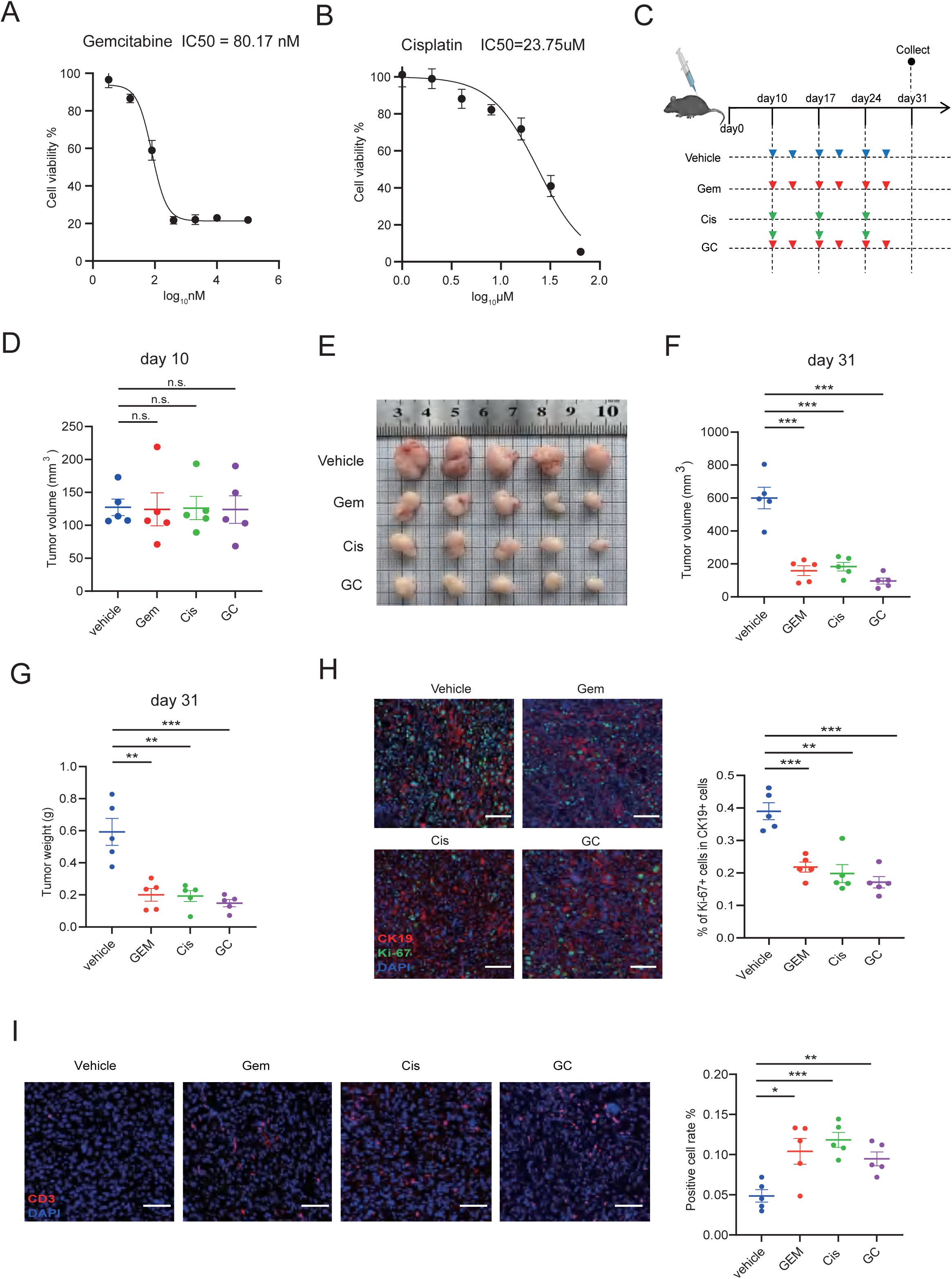
Drug sensitivity of mGBC1-ZH and subcutaneous tumor model. **(A)** Cell viability assay under gemcitabine treatment for 72 h at indicated doses. **(B)** Cell viability assay under cisplatin treatment for 72 h at indicated doses. **(C)** Workflow for the treatment of subcutaneous tumor-bearing mice. **(D)** The tumor volumes in vehicle- (n = 4), gemcitabine-treated (n = 4) and cisplatin-treated (n = 4) groups on day 10 after subcutaneous cell injection. n.s., not significant. **(E-G)** The tumors **(E)**, tumor volume **(F)** and tumor weight **(G)** of each group on day 30 after subcutaneous cell injection. *p < 0.05, **p <0.01, n.s., not significant. **(H)** Representative images and quantification of Ki-67+ proliferating cells in CK19+ epithelial cells in each group. Scale bars: 100 μm. **(I)** Representative images and quantification of CD3+ cells in each group. Scale bars: 100 μm

## Discussion

An ideal GBC mouse model should possess characteristics such as mimic human GBC genetic background and with a intact immune system. However, most of the currently available GBC mouse model are immune-deficient. In this study, by establishing a novel mouse GBC cell line harboring defined genetic drivers, we developed reproducible subcutaneous and orthotopic transplant gallbladder cancer models in syngeneic mice, addressing a significant gap in the field. Previously, only one mouse GBC cell line, A2, was reported (28). While A2 was derived from a transgenic mouse overexpressing ErbB2 in biliary epithelium under the control of bovine keratin 5 promoter, which developed tumors in both gallbladder and bile duct (29), further investigation using this cell line has been limited. An alternative in vivo model utilized an inducible Cre-based approach to simultaneously delete *PTEN* and activate *KRAS* within biliary epithelium, resulting in tumor development throughout the whole biliary system (30). By contrast, our model achieves superior tissue specificity through direct isolation of gallbladder tissue from adult mice.

A recent study by Shingo Kato et al. established an organoid-derived GBC mouse model through Cre-mediated recombination and CRISPR/Cas9 editing to activate *Kras* and delete *Trp53* (31). While their two-step implantation protocol successfully generated tumors on the gallbladder serosal surface in syngeneic mice, this model partially recapitulated the molecular features of Trp53-deficient cancers, as whole-genome sequencing revealed only focal copy number variations without detectable alterations in major cancer driver genes. In contrast, our development of the mGBC1-ZH cell line demonstrated pronounced genomic instability characterized by widespread chromosomal copy number alterations, thereby establishing a more representative murine model that faithfully mimics *Trp53* loss-associated carcinogenesis.

*Kras* activation and *Trp53* deletion mutations are the most prevalent genetic alterations observed in human GBC and known to be crucial for early disease development (32). Our transcriptomic analysis revealed significant alterations in gene expression during model development, with notable activation of PI3K-Akt and MAPK signaling pathways, consistent with human GBC molecular profiles. Genes exhibiting early expression changes at P7 represent potential drivers of tumor initiation and early progression. Further investigation on these genes may yield novel insights into GBC pathogenesis.Importantly, similar transcriptome between mGBC1-ZH and human GBC was observed through comparison with established human biliary tract cancers cell lines and single-cell sequencing data.Together, these results suggested that our model faithfully captured the main molecular characteristic of human GBC, thus may help elucidate the disease mechanisms and advancing our understanding of this heterogeneous disease.

We established syngeneic mouse models of GBC using mGBC1-ZH cells, generating both subcutaneous and orthotopic tumors in immunocompetent hosts. This approach overcomes key limitations of traditional xenograft models (4) by preserving an intact immune system. Histological characterization revealed T and B cell infiltration localized primarily to the tumor margins, indicating a “cold” immune microenvironment. Notably, we identified stromal cell-composed immune barrier structures in our orthotopic tumor, similar to those reported in hepatocellular carcinoma (18). This phenomenon parallel recent single-cell transcriptomic analyses of human GBC that revealed a complex stromal ecosystem with stromal-immune cell exclusivity (33). Investigating GBC immune evasion mechanisms using our novel model will provide novel insights in the future.

Using the established mGBC1-ZH cell line, we explored the role of CXCL5 in GBC tumorigenesis. CXCL5 is a classical chemokine known to play important roles in the development and progression of multiple cancer types (23,34). In our study, we demonstrated that CXCL5 promotes proliferation, metastasis, and invasion in both OCUG-1 and mGBC1-ZH cell lines. Besides these roles, as a member of the chemokine family, CXCL5 was identified as an inflammatory mediator with powerful role in neutrophil chemotaxis, and induced resistance to immunotherapy (35). Using our syngeneic GBC mouse model, we found that Cxcl5 promoted neutrophil infiltration and tumor malignancy in vivo. Recently, a multi-omics analysis of gallbladder cancerrevealed that tumoral CXCL5 expression is upregulated by AREG through the EGFR-pERK-EGR1 axis(36).Though they focus on the function of AREG, their work indicated that CXCL5 promotes neutrophil infiltration in metastatic tumors and contributes to resistance to immunotherapy. Collectively, these findings indicated that CXCL5 was a pivotal promoter of GBC progression, highlighting its potential as a therapeutic target and biomarker.

Gemcitabine plus cisplatin (GC) has been the standard first-line chemotherapy for GBC for over a decade (37). Our syngeneic model demonstrated sensitivity to both agents, with response patterns consistent with clinical observations. This pharmacological validation establishes our model as a reliable preclinical platform for therapeutic evaluation. While our current study focused on conventional chemotherapeutics, this immunocompetent model is particularly well-suited for investigating emerging immunotherapeutic approaches.

In conclusion, the mGBC1-ZH cell line and derived syngeneic mouse model represent an ideal experimental platform for GBC research. By overcoming the limitations of existing models, this system enables comprehensive investigation of GBC pathogenesis, the tumor immune microenvironment, and novel treatment approaches, thereby advancing both basic and translational research in GBC.

## Abbreviations

GBC: Gallbladder cancer
PDXs: patient-derived xenografts
3D: three-dimensional
EMT: epithelial-mesenchymal transition
TIB: tumor immune barrier
STR: Short Tandem Repeat
WES: whole-exome sequencing
PDAC: pancreatic ductal adenocarcinoma
GEM: gemcitabine
CIS: cisplatin
IC50: half-maximal inhibitory concentration

## Funding

This work was supported by funding from the National Natural Science Foundation of China (32300484 and 82373311), the Science and Technology Commission of Shanghai Municipality (22ZR1411700 and 21JC1401200), and Xuhui Academy-Local Collaboration Initiative in Life and Health (25XHYD-02).

## Conflict of interest

A patent application has been filed based on the findings in this paper (Houbao Liu et al. CN202511038362.4).

## Authors’ contributions

Q.W., Y.L., L.W., Y.W., and H.L. contributed to study concept and design. Q.W. and X.N. completed the acquisition of data with assistance from F.X., W.M., R.Q., J.M., B.L., S.X., C.W., L.N., and X.B. Analysis and interpretation of data were performed by Q.W., W.M.,and D.Z. Q.W. and Y.L. wrote the manuscript with assistance from all authors. Q.W., L.W., Y.W. and H.L. supervised the work.

## Data Availability Statement

The data that support the findings of this study are available from the corresponding author upon reasonable request.

## Notes

### Summary of Updates

RNA-seq analysis of the shNT and shCxcl5 cells of mGBC1-ZH. Tumor growth comparison of CT and Cxcl5-OE cells in C57BL/6J mouse, quantification of H&E, immunohistochemical for Ki-67, Ly6G, and MPO of subcutaneous tumors.

