## Supplementary Figure 1 for "A Syngeneic Immunocompetent Mouse Model of Gallbladder Cancer Reveals Tumorigenesis and Therapeutic Response"

A

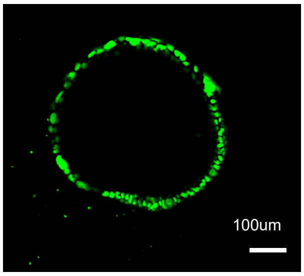


B

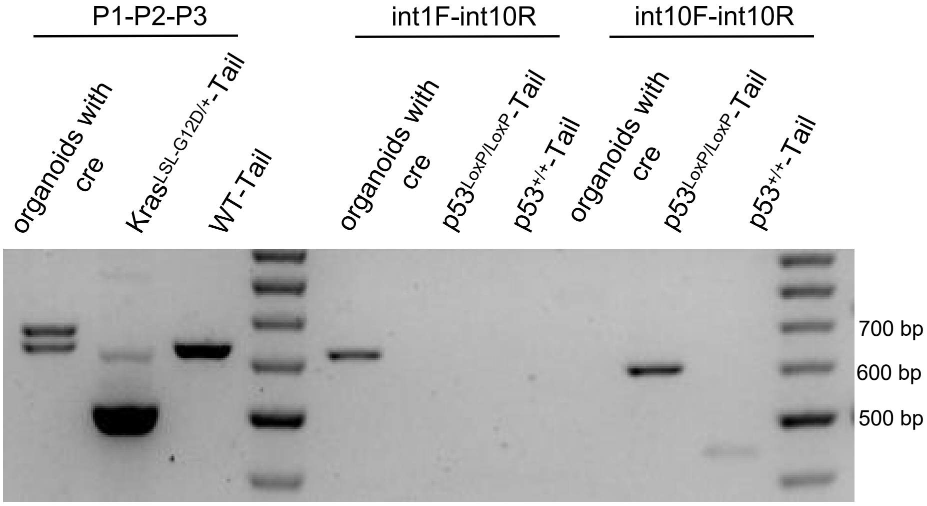


| | P53 | | Kras |
|---|---|---|---|
| | Primer int1-int10 | Primer int10-int10 | Primer P1-P2-P3 |
| CRE-recombined | 612bp | - | 650bp、622bp |
| p53loxp/loxp | - | 584bp | 622bp、500bp |
| WT | - | 431bp | 622bp |

**Fig. S1. The efficiency of Cre-mediated recombination. (A)** Representative image of organoids after introduction with the lentivirus vector with Cre and a tracking marker EGFP. Scale bars: 100 μm. **(B)** PCR analysis of Cre-mediated recombination. The length of PCR proudcts corresponding to different conditions were listed in the table. Primers were listed in Supplementary Table S4.
