## Supplementary Figure 2 for "A Syngeneic Immunocompetent Mouse Model of Gallbladder Cancer Reveals Tumorigenesis and Therapeutic Response"

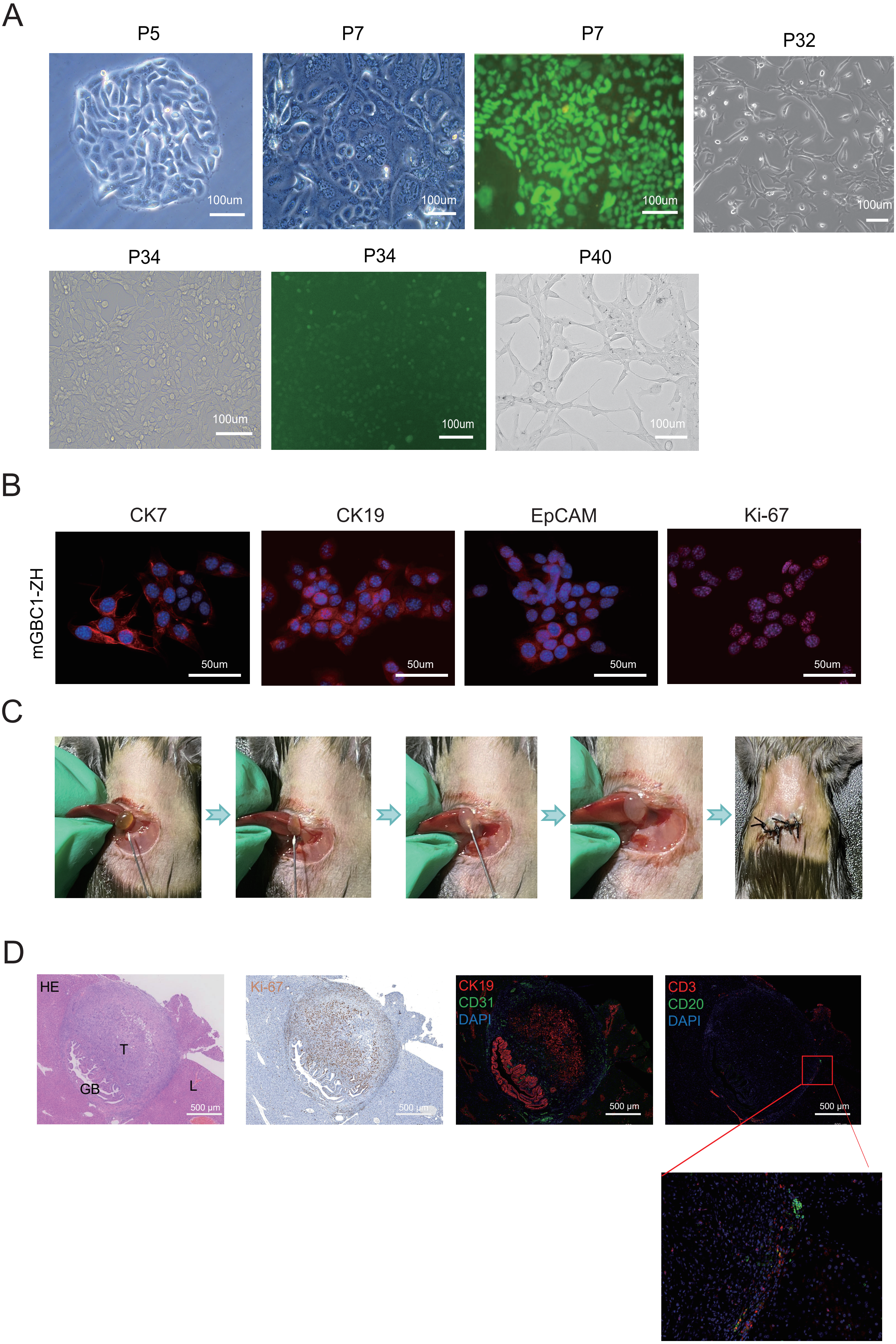

**Fig. S2. The generation of mGBC1-ZH cell line and orthotopic tumor. (A)** Representative bright field images and GFP expression of cells at various passage numbers. **(B)** Immunofluorescence for CK7, CK19, EpCAM and Ki-67 of mGBC1-ZH. **(C)** Schematic of the strategy for generation the orthotopic gallbladder cancer model. The gallbladder was punctured to extrude bile, cells resuspened in matrigel was injected to the gallbladder. **(D)** H&E, immunohistochemical for Ki-67, immunofluorescence for CK19, CD31, CD3 and CD20 of a orthotopic tumor. The tumor was surrounded by stromal cells.
