## Supplementary Figure 3 for "A Syngeneic Immunocompetent Mouse Model of Gallbladder Cancer Reveals Tumorigenesis and Therapeutic Response"

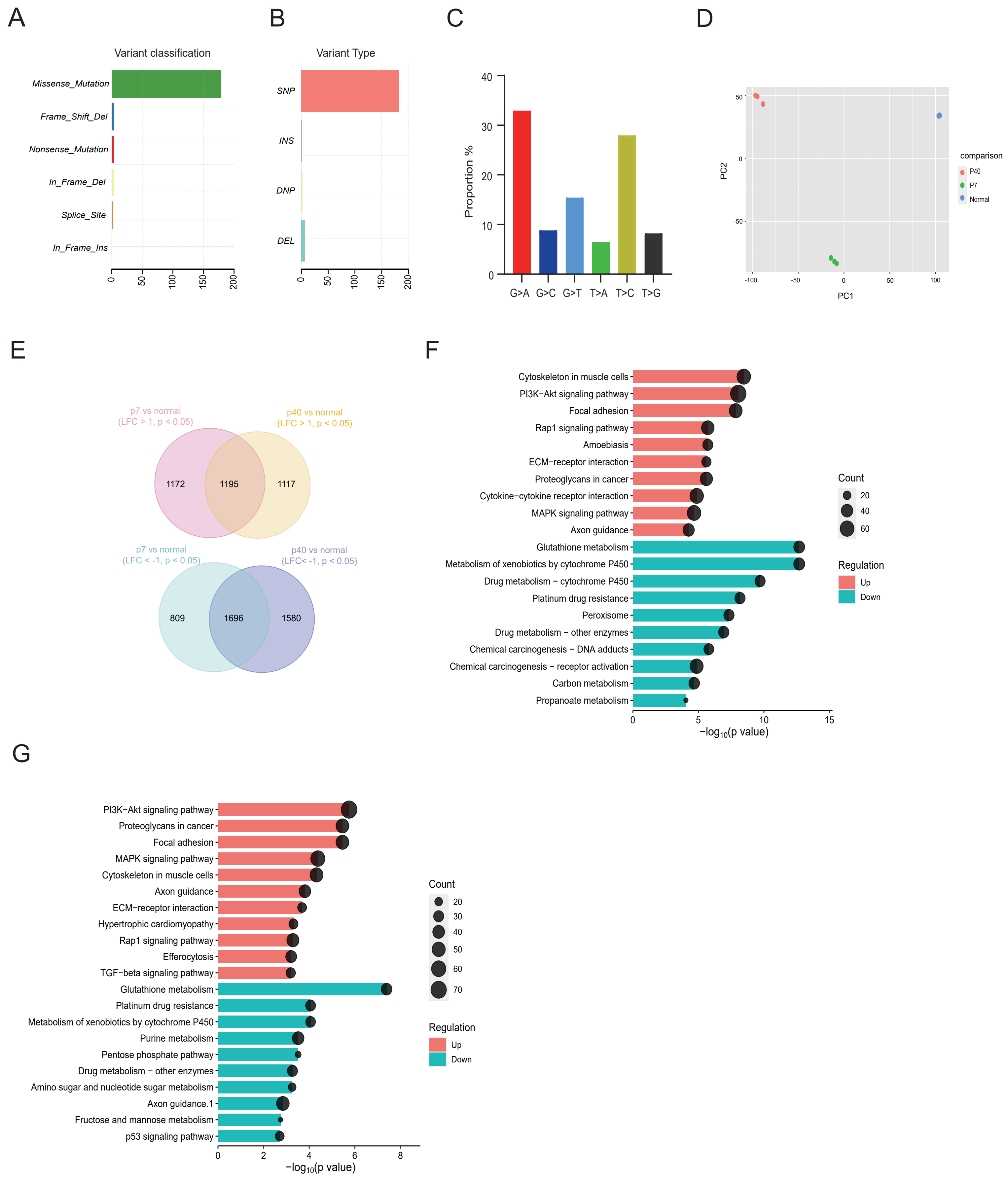

**Fig. S3. Transcriptome and genetic characteristics of mGBC1-ZH. (A)** Bar plot showing the number of different classes of variants identified in mGBC1-ZH. **(B)** Bar plot showing the number of different classes of variants types identified in mGBC1-ZH. **(C)** The distribution of the six types of base substitutions detected in mGBC1-ZH. **(D)** PCA map for gene expression profiles of Normal, P7 and P40. **(E)** Venn plot of overlapped differential expression genes identified between P7 and Normal, and P40 and Normal. **(F)** Pathway enrichment analysis of the differential expression genes identified between P7 and Normal using the KEGG database. **(G)** Pathway enrichment analysis of the differential expression genes identified between P40 and Normal using the KEGG database.
