## Supplementary Figure 4 for "A Syngeneic Immunocompetent Mouse Model of Gallbladder Cancer Reveals Tumorigenesis and Therapeutic Response"

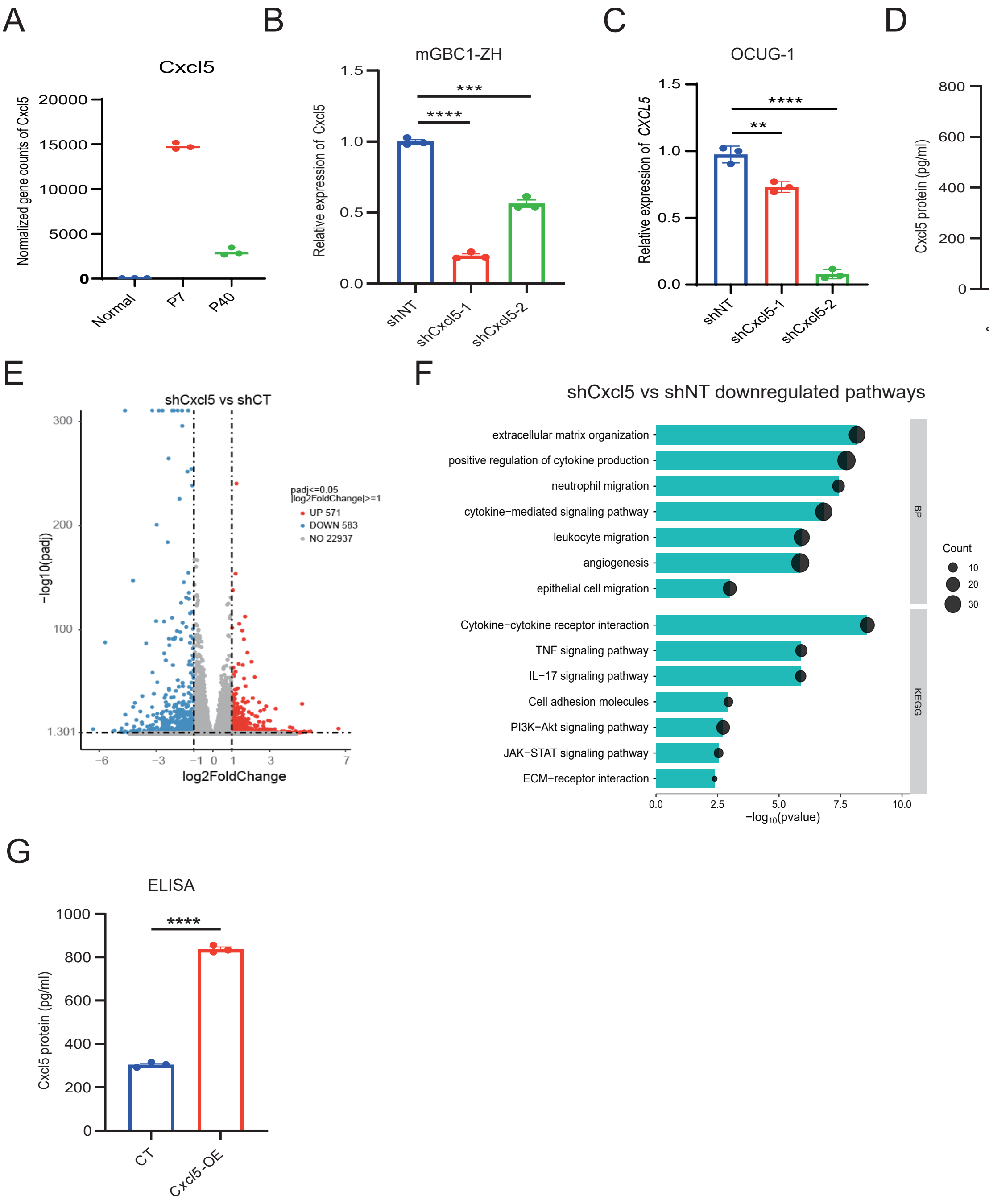

**Fig. S4. Validation of CXCL5 stable cell lines. (A)** The normalized gene counts of *Cxcl5* in Normal, P7 and P40 cells. **(B)** qPCR analysis of Cxcl5 expression in shNT and shCxcl5 cells of mGBC1-ZH. **(C)** qPCR analysis of Cxcl5 expression in shNT and shCxcl5 cells of OCUG-1. **(D)** ELISA analysis of Cxcl5 concentration in the culture supernatants in shNT and shCxcl5 cells of mGBC1-ZH. **(E)** Volcano plot of the differential expression genes identified between shNT and shCxcl5 cells of mGBC1-ZH. **(F)** Go and KEGG enrichment of the down-regulated genes in shCxcl5 compared to shNT. **(G)** qPCR analysis of Cxcl5 expression in CT and Cxcl5-OE cells of mGBC1-ZH. **(H)** ELISA analysis of Cxcl5 concentration in the culture supernatants in CT and Cxcl5-OE cells of mGBC1-ZH.
